# Antigenicity and receptor affinity of SARS-CoV-2 BA.2.86 spike

**DOI:** 10.1101/2023.09.24.559214

**Authors:** Qian Wang, Yicheng Guo, Liyuan Liu, Logan T. Schwanz, Zhiteng Li, Jerren Ho, Richard M. Zhang, Sho Iketani, Jian Yu, Yiming Huang, Yiming Qu, Riccardo Valdez, Adam S. Lauring, Aubree Gordon, Harris H. Wang, Lihong Liu, David D. Ho

## Abstract

Although the COVID-19 pandemic has officially ended^1^, SARS-CoV-2 continues to spread and evolve. Recent infections have been dominated by XBB.1.5 and EG.5.1 subvariants^2^. A new subvariant designated BA.2.86 has just emerged, spreading to 21 countries in 5 continents^3^. This virus contains 34 spike mutations compared to its BA.2 predecessor, thereby raising concerns about its propensity to evade existing antibodies. We examined its antigenicity using human sera and monoclonal antibodies (mAbs). Reassuringly, BA.2.86 was not more resistant to human sera than XBB.1.5 and EG.5.1, indicating that the new subvariant would not have a growth advantage in this regard. Importantly, sera from patients who had XBB breakthrough infection exhibited robust neutralizing activity against all viruses tested, suggesting that upcoming XBB.1.5 monovalent vaccines could confer added protection. The finding that the longer genetic distance of BA.2.86 did not yield a larger antigenic distance was partially explained by the mAb data. While BA.2.86 showed greater resistance to mAbs to subdomain 1 (SD1) and receptor-binding domain (RBD) class 2 and 3 epitopes, it was more sensitive to mAbs to class 1 and 4/1 epitopes in the “inner face” of RBD that is exposed only when this domain is in the “up” position. We also identified six new spike mutations that mediate antibody resistance, including E554K that threatens SD1 mAbs in clinical development. The BA.2.86 spike also had a remarkably high receptor affinity. The ultimate trajectory of this new SARS-CoV-2 variant will soon be revealed by continuing surveillance, but its worldwide spread is worrisome.

## INTRODUCTION

A highly mutated severe acute respiratory syndrome coronavirus 2 (SARS-CoV-2) Omicron subvariant, designated BA.2.86, was first reported only a few weeks ago, and it is genetically distinct from the prevailing viruses in the XBB sublineage^2,4-6^. The genetic distance to its predecessor, BA.2, is equivalent to that between BA.1 and the Delta variant (**Figure 1A**), raising the same antibody evasion concerns when the first Omicron variant emerged in late 2021. Over 190 sequences of BA.2.86 has been found in 21 countries^3^ already despite limited surveillance nowadays. A recent outbreak due to the new subvariant in a nursing facility in England with high attack rate among residents and staff shows BA.2.86 is readily transmissible^7^. At present, there is little clinical evidence to address its pathogenicity.

**Figure 1.**
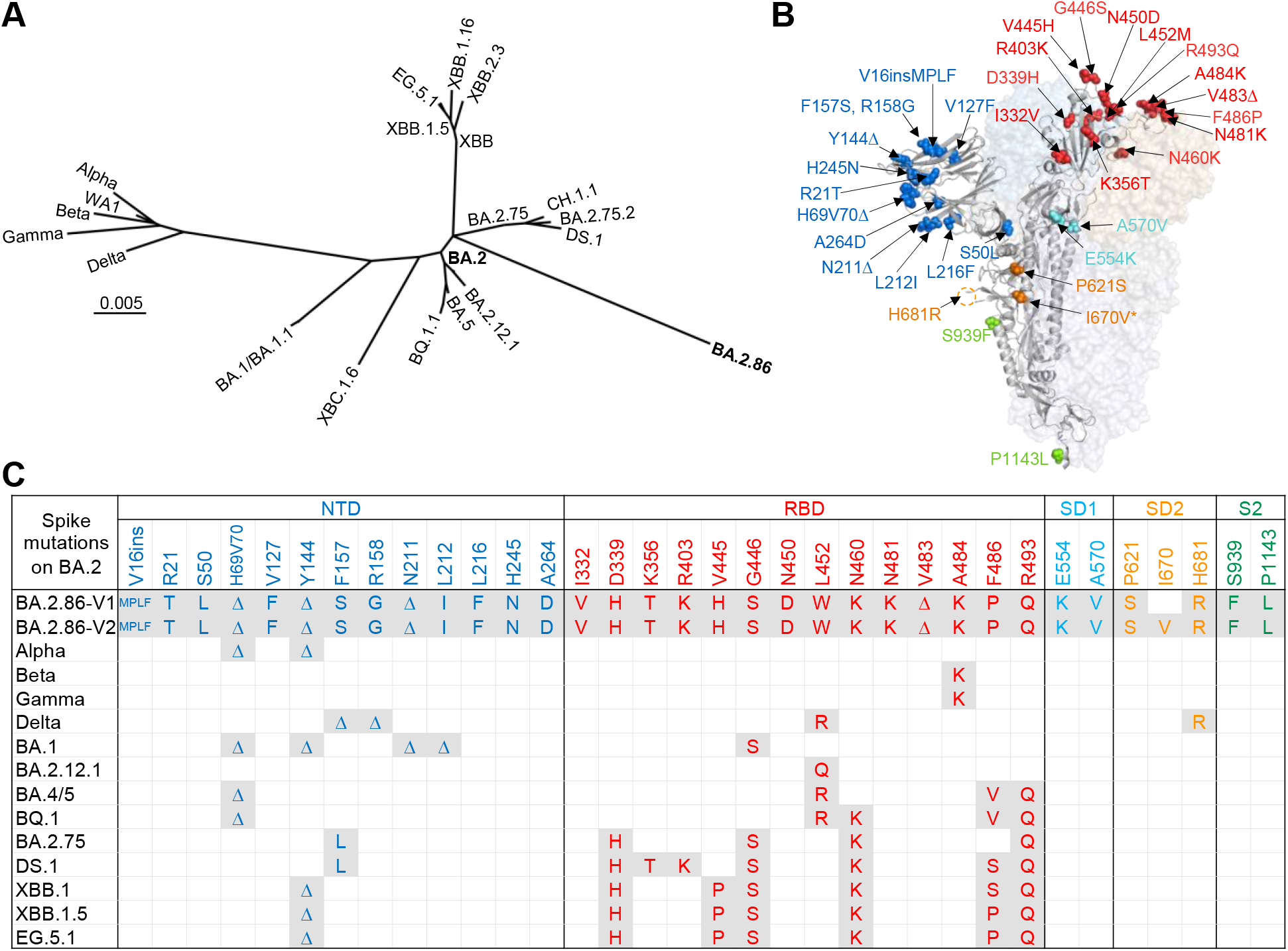
Divergence of BA.2.86 spike sequence from major SARS-CoV-2 variants. A. Phylogenetic tree of SARS-CoV-2 variants based on spike sequences. B. Location of mutations detected in BA.2.86 spike, relative to its ancestral BA.2. The red, blue, cyan, orange, and green mutations are in RBD, NTD, SD1, SD2, and S2, respectively. The orange circle indicates the H681R mutation located proximal to the furin cleavage site. I670V denoted by an asterisk, since it is found in only a minority of BA.2.86 spikes (BA.2.86-V2); the dominant form does not have this mutation (BA.2.86-V1). ins, insertion; Δ, deletion. C. Spike mutations found in BA.2.86 and other SARS-CoV-2 variants compared with BA.2.

Compared with the spike of BA.2, BA.2.86 possesses 34 additional mutations, including 13 mutations in the N-terminal domain (NTD), 14 in RBD, 2 in SD1, 3 in the subdomain 2 (SD2), and 2 in the S2 region (**Figures 1B and 1C**). Mutations H69V70 deletion (H69V70Δ), Y144 deletion (Y144Δ), G446S, N460K, F486P, and R493Q have been identified previously^5,6^,8,9, but mutations V445H, N450D, N481K, V483 deletion (V483Δ), and E554K have been seldom observed in circulating viruses (**Figure 1C**). This extensive array of spike mutations in BA.2.86 is alarming because of the heightened potential for the virus to evade serum antibodies elicited by prior infections and/or vaccinations or mAbs intended for clinical use. The present study addresses this concern by characterizing the antigenicity of BA.2.86 spike using multiple collections of human sera and a large panel of mAbs.

## RESULTS

### Sequence variation

The initial analysis of available BA.2.86 spike sequences was challenging due to sequence variations and uncertainties. A four amino-acid insertion after the V16 residue (V16insMPLF) was observed in a majority of reported sequences, while some were ambiguous because of low sequencing quality spanning this region (**Extended Data Figure 1**). We therefore made the determination that V16insMPLF should be included in our spike construct. Another variation is the presence or absence of the I670V mutation. Before it was recognized that most BA.2.86 strains do not contain this mutation (**Extended Data Figure 1**), we already synthesized both spike genes by methods previously described^4,10^: BA.2.86-V1 being the dominant form and BA.2.86-V2 being the minor form (**Figure 1C**).

### Serum neutralization

To assess the antigenicity of the BA.2.86 spike, we constructed vesicular stomatitis virus (VSV) pseudotyped viruses using both versions of the spike gene, as well as BA.2, XBB.1.5, and EG.5.1 pseudoviruses for comparison. These pseudoviruses were then subjected to neutralization studies using serum samples from three distinct clinical cohorts. The first cohort consisted of healthy individuals who received three doses of monovalent mRNA vaccines followed by two doses of BA.5 bivalent mRNA vaccines (referred to as “3 shots monovalent + 2 shots bivalent”). The other two cohorts included patients who experienced a breakthrough infection caused by BA.2 (labeled as “BA.2 breakthrough”) or XBB (labeled as “XBB breakthrough”) after multiple vaccinations. More details on the clinical samples can be found in **Extended Data Table 1**.

The serum neutralization results and comparative analyses are shown in **Figure 2A** and **Figure 2B**, respectively. BA.2.86-V1 and BA.2.86-V2 displayed comparable neutralization ID_50_ (50% inhibitory dilution) titers across all three cohorts, indicating that the I670V mutation has no appreciable antigenic impact. Among the variants tested, BA.2 was most sensitive to neutralization by sera from all three cohorts. Surprisingly, BA.2.86 was not the most resistant; EG.5.1 was instead. In fact, compared to XBB.1.5 and EG.5.1, BA.2.86 was 1.5- and 2.0-fold, respectively, more sensitive to neutralization by sera from the “3 shots monovalent + 2 shots bivalent” cohort. BA.2.86 was also more sensitive to neutralization by sera from the “BA.2 breakthrough” cohort than EG.5.1 by 1.9-fold. BA.2.86, XBB.1.5, and EG.5.1 were similarly sensitive to neutralization by sera from the “XBB breakthrough” cohort; notably, the serum ID_50_ titers were quite robust, ranging from 729 to 879. This result suggests that exposure to the spike of XBB.1.5 could lead to an effective antibody response against the current circulating SARS-CoV-2 variants, an inference that bodes well for the upcoming XBB.1.5 monovalent vaccines.

**Figure 2.**
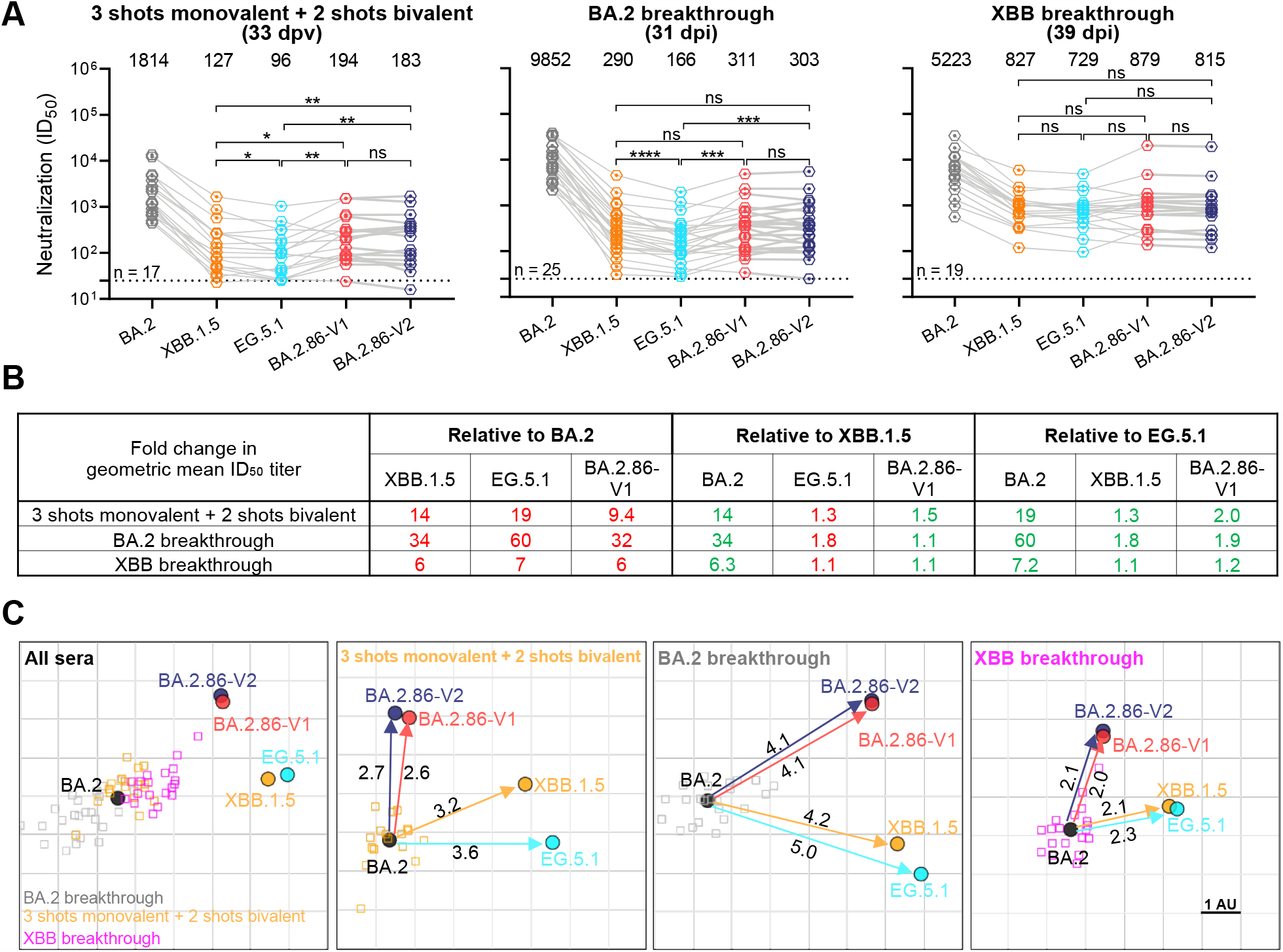
Serum neutralization of BA.2.86 compared with BA.2, XBB.1.5, and EG.5.1. A. Neutralizing ID_50_ titers of serum samples from “3 shots monovalent + 2 shots bivalent”, “BA.2 breakthrough” and “XBB breakthrough” cohorts against the indicated SARS-CoV-2 variants. The geometric mean ID_50_ titers (GMT) are presented above symbols. The neutralization assay limit of detection (dotted line) is 25. Statistical analyses were performed by employing Wilcoxon matched-pairs signed-rank tests. ns, not significant; \**p* < 0.05; \*\**p* < 0.01; \*\*\**p* < 0.001; \*\*\*\**p* < 0.0001. n, sample size. dpv, days post last vaccination; dpi, days post infection. BA.2.86-V2 carries an I670V mutation compared to the dominant version of BA.2.86 (BA.2.86-V1). B. Fold changes in GMT relative to BA.2, XBB.1.5, and EG.5.1, with resistance colored red and sensitization colored green. C. Antigenic map generated using neutralization data from panel A. BA.2 represents the central reference for all serum cohorts, with the antigenic distances calculated by the average divergence from each variant. One antigenic unit (AU) represents an approximately 2-fold change in ID_50_ titer.

The serum neutralization data were then used to generate antigenic maps to graphically show the antigenic relationships between BA.2.86 and the other Omicron subvariants tested (**Figure 2C**). The scientific conclusions are obviously the same as those already stated, but such a display allows easier visualization of the overall findings.

### Neutralization by mAbs

To understand the antibody evasion properties of BA.2.86 in greater detail, we evaluated the susceptibility of the dominant form, BA.2.86-V1, to neutralization by a panel of 25 mAbs that retained activity against BA.2. XBB.1.5 and EG.5.1 were included as comparators. Among the mAbs, 20 target the four epitope classes in the RBD^11^, including S2K146^12^, BD57-0129^13^, BD56-1302^13^, DB56-1854^13^, Omi-3^14^, Omi-18^14^, BD-515^15^, Omi-42^14^, COV2-2196 (tixagevimab)^16^, XGv347^17^, ZCB11^18^, XGv051^17^, A19-46.1^19^, S309 (sotrovimab)^20^, COV2-2130 (cilgavimab)^16^, LY-CoV1404 (bebtelovimab)^21^, Beta-54^22^, BD55-4637^13^, SA55^23^, and 10-40^24^. The other 5 mAbs were C1520^25^ targeting the NTD, C1717^25^ targeting both the NTD and subdomain 2 (NTD-SD2), and 3 SD1-directed monoclonals, including S3H3^26^ and two antibodies (ADARC1 and ADARC2) that we have been characterizing (our unpublished results). The raw IC_50_ (50% inhibitory concentration) values are shown in **Extended Data Table 2**, and fold changes in IC_50_ titers relative to BA.2 are summarized in **Figure 3A**.

**Figure 3.**
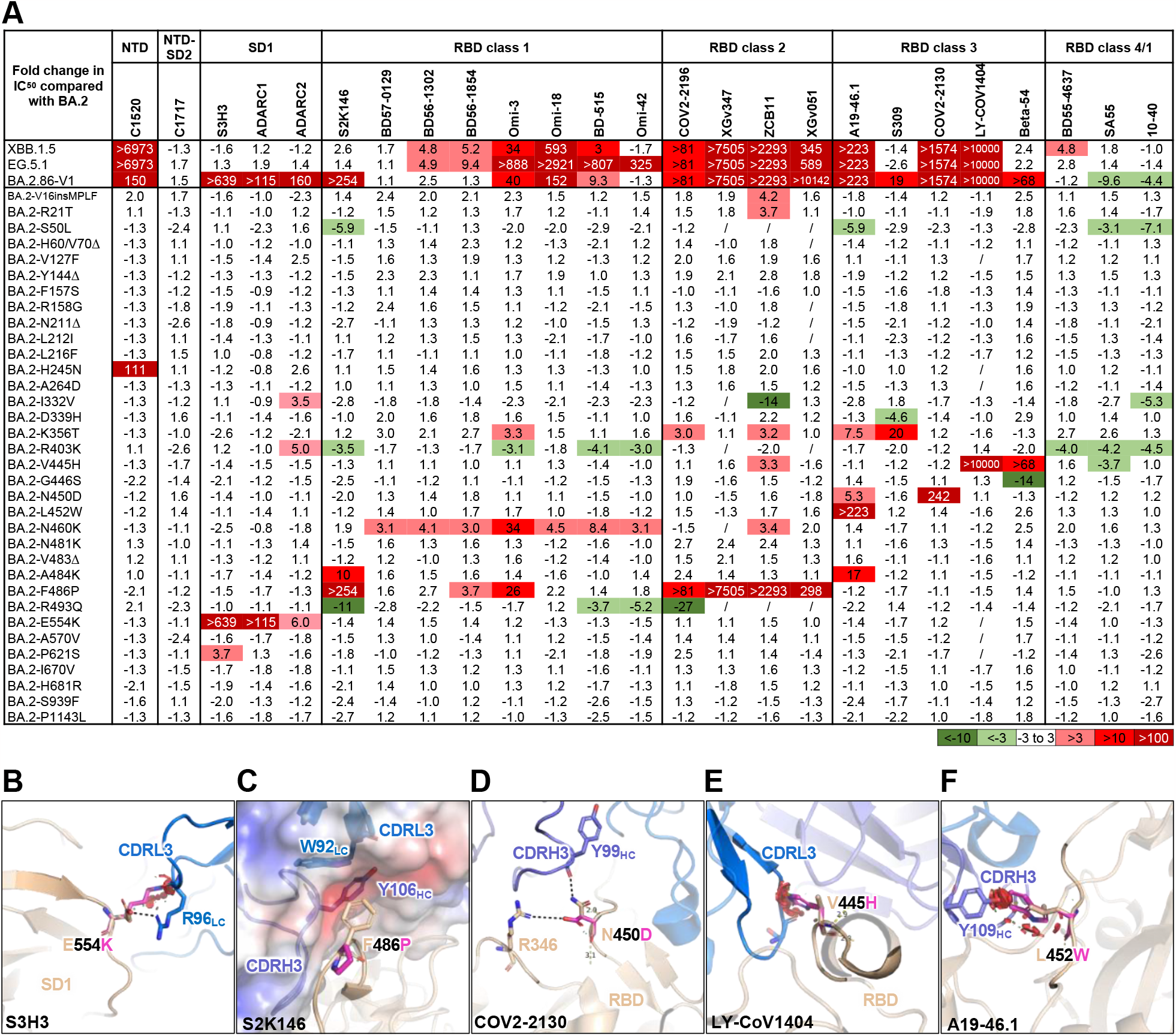
Neutralization of BA.2.86 and its point mutants in BA.2 by a panel of mAbs. A. Fold changes in IC_50_ values of XBB.1.5, EG.5.1, BA.2.86-V1, and point mutants relative to BA.2, with resistance colored red and sensitization colored green. “/”, fold change not available as the IC_50_ value was below the limit of detection (< 0.001 μg/mL). B-F. Structural modeling of how single mutations affect S3H3 (b), S2K146 (c), COV2-2130 (d), LY-CoV1404 (e), A19-46.1 (f) neutralization. Dashed lines indicate salt bridges or hydrogen bonds. Red plates indicate steric hindrance. The surfaces are colored according to the electrostatic potential of mAb S2K146.

Our results revealed that BA.2.86 was completely or substantially resistant to neutralization by mAbs to NTD, SD1, and RBD class 2 and class 3 epitopes, and the extent of its evasion from such antibodies appeared larger than those exhibited by XBB.1.5 and EG.5.1. In particular, BA.2.86 showed greater resistance to RBD class 2 mAb XGv051 and class 3 mAbs S309 and Beta-54, while escaping almost completely from SD1 mAbs that could neutralize both XBB.1.5 and EG.5.1. Unexpectedly, BA.2.86 was substantially more sensitive to neutralization than EG.5.1 by a majority mAbs to class 1 and class 4/1 epitopes on the ‘inner face’ of RBD that are only revealed when this domain is in the “up” position^11,27.^ This observation suggests that the RBD of BA.2.86 may be more exposed and accessible to certain antibodies. Overall, the opposing effects of different mutations on different classes of antibodies also explain, in part, why the longer genetic distance did not translate into a larger antigenic distance for BA.2.86.

To elucidate the impact of each BA.2.86 spike mutations on its antigenicity, we synthesized the gene for each of the 34 point mutants in the background of BA.2 and then constructed the corresponding pseudoviruses for neutralization studies using the same panel of mAbs (**Figure 3A**). The H245N mutation mediated resistance to the NTD antibody C1520. Significantly, the E554K mutation conferred evasion to all SD1-directed antibodies tested, which is in line with a report on a different SD1 mAb^28^. Structural modeling suggests that E554K removes the salt bridge formed between E554 and R96 in the CDRL3 region of mAb S3H3 and induces steric hindrance that disrupts antibody binding (**Figure 3B**). Mutations N460K and F486P, also shared by XBB.1.5 and EG.5.1, mediated resistance to some RBD class 1 and/or class 2 mAbs. Specifically, the N460K mutation, first observed in the BA.2.75 variant, disrupts a key hydrogen bond between the RBD and VH3-53-encoded class 1 antibodies^29^, but enhancing receptor affinity^30^ at the same time. The F486P mutation appears to reduce the hydrophobic interaction with the ACE2-mimicking antibody S2K146 (**Figure 3C)**, hence impairing its neutralization activity. The K356T mutation, also shared by the DS.1 variant, conferred broad resistance to a number of RBD class 1, class 2, and class 3 mAb, possibly due to steric hindrance caused by the introduction of an additional glycosylation site^8^. Several other RBD mutations, including V445H, N450D, L452W, and A484K compromised the neutralizing activity of some RBD class 3 mAbs. Structural modeling indicates that N450D could form an additional salt bridge with R346, thereby altering the local conformation and resulting in resistance to mAbs such as COV2-2130 (**Figure 3D)**. On the other hand, mutations V445H and L452W seem to introduce steric clashes with the CDRs of RBD class 3 mAbs LY-CoV1404 (**Figure 3E**) and A19-46.1 (**Figure 3F**), respectively. Importantly, we also found two new mutations (S50L and I332V) that conferred a degree of sensitization to neutralization by certain mAbs, along with two previously known mutations (R403K and R493Q) that were also sensitizing^8,9^,31 (**Figure 3A**). The rest of the new mutations that are unique to BA.2.86 showed only minor or no effect on its antigenicity as assessed by this panel of mAbs. In summary, a number of mutations in this new variant caused resistance to antibody neutralization, and several other mutations mediated an opposite effect, while the remainder were antigenically neutral.

### Receptor affinity

We also expanded our studies on the BA.2.86 spike by measuring its binding affinity to the viral receptor. The spike proteins of BA.2.86-V1 and BA.2.86-V2, along with those of BA.2, XBB.1.5, and EG.5.1 were first examined for binding to a dimeric human-ACE2-Fc protein by surface plasmon resonance (SPR) as we have previously reported^9^. XBB.1.5 and EG.5.1 spikes exhibited comparable affinities to ACE2, with K_D_ values of 1.34 nM and 1.21 nM, respectively (**Figure 4A**). These values represent only a modest increase in receptor binding affinity compared to the K_D_ value of the BA.2 spike (1.68 nM). In contrast, both versions of the BA.2.86 spikes showed a >2-fold increase in binding affinity, with similar K_D_ values of 0.54 nM and 0.60 nM, largely due to lower dissociation rates (K_d_).

**Figure 4.**
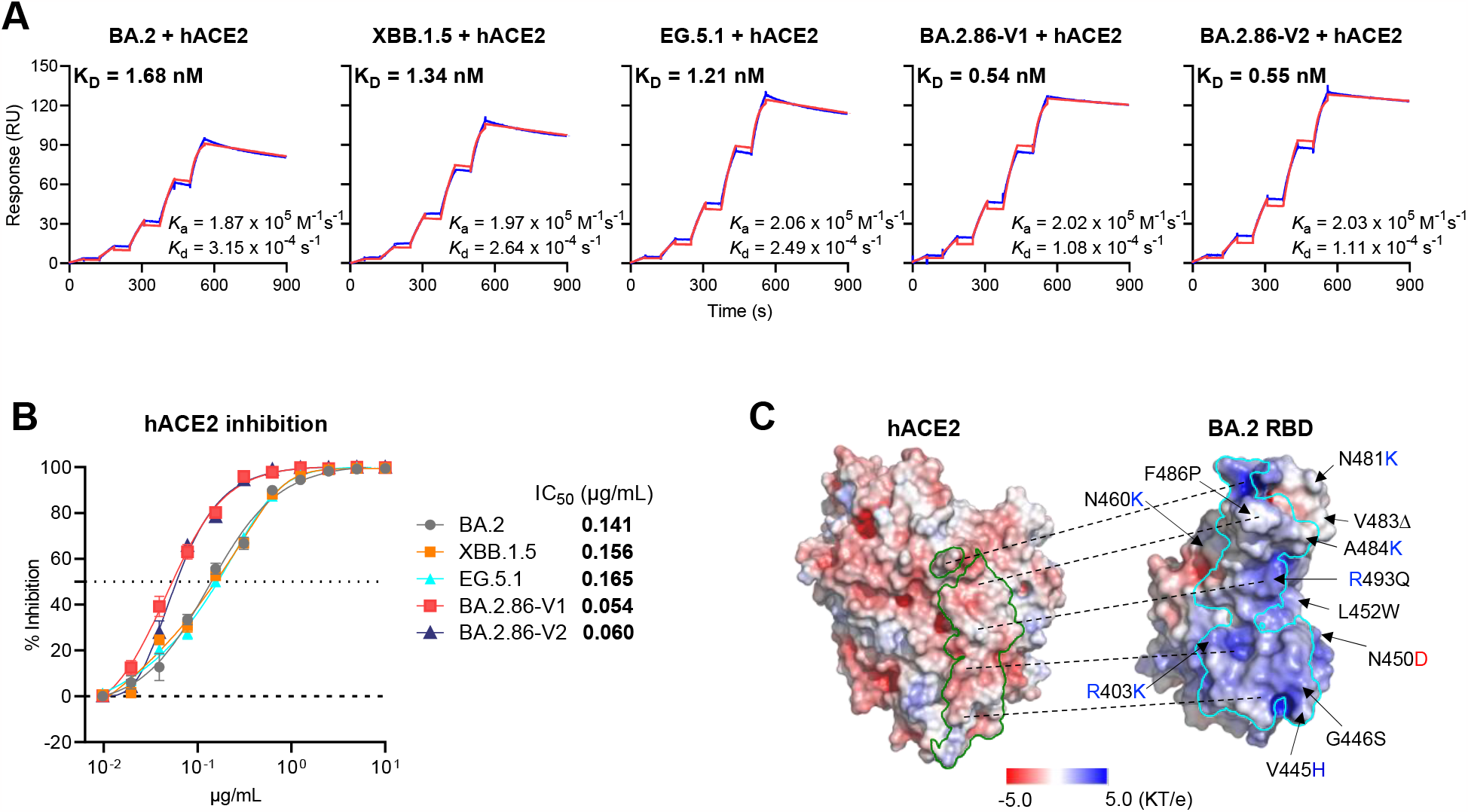
BA.2.86 exhibited stronger receptor affinity than BA.2, XBB.1.5 and EG.5.1. A. ACE2 receptor binding affinity of BA.2.86 spike, in comparison with spikes from BA.2, XBB.1.5, and EG.5.1 as tested by SPR. B. Susceptibility of two versions of BA.2.86 pseudoviruses to hACE2 inhibition, relative to that of BA.2, XBB.1.5, and EG.5.1. Data are shown as mean ± standard error of mean (SEM). C. Electrostatic potential of hACE2 and the BA.2 RBD, with arrows indicating the mutations identified in BA.2.86. The green and cyan boundaries delineate the footprints of the RBD and hACE2, respectively. The dashed lines showed the corresponding interaction surfaces between RBD and hACE2. Residues with positive and negative charges are colored as blue and red, respectively.

To corroborate these findings, we also evaluated the susceptibility of both BA.2.86 pseudoviruses to neutralization by the dimeric human-ACE2-Fc protein, in comparison to BA.2, XBB.1.5, and EG.5.1. In agreement with the SPR data, both versions of BA.2.86 were >2-fold more sensitive to ACE2 inhibition than XBB.1.5 and EG.5.1, as determined by their IC_50_ values (**Figure 4B**). A potential explanation for this heightened affinity may reside in the intrinsic charge properties of the two interacting molecules. The region of human ACE2 targeted by the RBD is negatively charged, while the Omicron RBD itself is positively charged^32^. The higher receptor binding affinity of the BA.2.86 spike might be attributed to the additional positive charges associated with mutations V445H, N460K, N481K and A484K (**Figure 4C**). Only the N460K mutation is shared with the spikes from XBB.1.5 and EG.5.1.

## Discussion

SARS-CoV-2 variant BA.2.86 emerged only recently, but it has raised alarm because of the extensive array of mutations in its spike protein. Current concerns about its antibody evasiveness are reminiscent of those when the first Omicron appeared in late 2021. We have therefore undertaken a thorough antigenic characterization of BA.2.86, and our findings have important clinical and scientific implications.

On the clinical front, our data showed that, compared to the currently dominant subvariants XBB.1.5 and EG.5.1, BA.2.86 did not exhibit greater resistance to neutralization by human sera from three different cohorts in the United States (**Figure 2A**). In fact, it was slightly but appreciably more sensitive to serum neutralization than EG.5.1 (**Figure 2B**). Our results are in concordance with findings posted by Lasrado et al^33^ from the US as well as results posted by Khan et al^34^ from South Africa, but in contrast with those posted by Yang et al from China^35^, Uriu et al from Japan^36^, and Sheward et al from Sweden^37^, who found BA.2.86 to be slightly more resistant to antibodies in human sera. The discrepancy with the latter reports could be due to differences in the histories of exposures to SARS-CoV-2 infection and/or vaccination. Going forward, it will be important to understand the basis of the observed discrepancies, because relatively greater resistance to antibody neutralization could confer an advantage for the new variant to grow in the population.

Another clinical ramification of our findings is that the upcoming XBB.1.5 monovalent vaccines are likely to elicit an adequate antibody response to not only BA.2.86 but also the currently dominant subvariants XBB.1.5 and EG.5.1. This reassuring conclusion is inferred from our results showing that sera from the “XBB breakthrough” cohort exhibited robust neutralization titers against all viral variants tested (**Figure 2A**), but more importantly it is now confirmed by results just posted by Moderna on its monovalent XBB.1.5 mRNA vaccine^38^. They too noted that BA.2.86 was not more resistant to antibody neutralization than XBB.1.5 and EG.5.1.

A third clinically relevant result is the loss of neutralizing activity for all of the SD1-directed mAbs we tested against BA.2.86. One previous study highlighted that SD1 antibodies are rarely induced by infection or vaccination^28^, raising the specter that such antibodies could possibly maintain its neutralizing activity durably in the face of continuing SARS-CoV-2 evolution and become ideal candidates for clinical development. Regrettably, BA.2.86 by making the E554K mutation (**Figure 3A**) has dashed any such hope.

Our detailed studies on a panel of mAbs have also yielded important scientific insights on the evolutionary pathways taken by SARS-CoV-2. We have previously noted that Omicron subvariant XBC.1.6 exhibited a longer genetic distance from the ancestral virus than EG.5.1, and yet it was more sensitive to antibody neutralization than EG.5.1^2^. That observation remained unexplained, but now a parallel situation has arisen with BA.2.86 that could be partially explained by our new findings. While BA.2.86 showed greater resistance to mAbs to SD1 and RBD class 2 and 3 epitopes, it was more sensitive to mAbs to RBD class 1 and 4/1 epitopes (**Figure 3A**). Moreover, a number of its mutations (e.g., K356T, V445H, N450D, E460K, F486P, and E554K) conferred antibody resistance, but their neutralizing effects are offset by other mutations (e.g., S50L, I332V, R403K, and R493Q) that conferred antibody sensitization.

Another scientific implication of our results is that the RBD of BA.2.86 is likely to be more exposed than the RBD of XBB.1.5 or EG.5.1. This conclusion is inferred from the above observation that the new variant is more sensitive than XBB.1.5 or EG.5.1 to neutralization by class 1 and 4/1 mAbs, which target the “inner face” of RBD only when this domain is in the “up” position. Since receptor binding also occurs when the RBD is “up”, this conclusion is in line with the finding that the spike of BA.2.86 has a >2-fold higher affinity for the viral receptor compared to the spike of XBB.1.5 or EG.5.1 (**Figures 4A and 4B**). In fact, BA.2.86 spike has one of the highest receptor affinities we have measured, together with the spikes of some of the viruses in the BA.2.75 sublineage^8^ but the K_D_ is undoubtedly determined by additional properties including the electrostatic charge of the RBD (**Figures 4C**).

We have witnessed, almost in real time, the evolution of SARS-CoV-2 over the past three years. Studies on the successive waves of viral variants and subvariants have taught us that this virus is constantly mutating to evade pressure exerted by antibodies in human sera. Given the extent of herd immunity today, only the most antibody resistant forms will have a growth advantage and become dominant. At the same time, the spikes of recently dominant variants all possess high receptor affinity, which is one measure of viral fitness. The trajectory of BA.2.86 ahead will be determined by the characteristics described herein as well as by viral mutations beyond spike and yet to be defined host factors. However, the fact that this emerging variant has already spread to so many different countries scattered around the world would suggest that it must be quite fit, and that continuing surveillance is imperative.

## ACKNOWLEDGEMENTS

This study was supported by funding from the NIH SARS-CoV-2 Assessment of Viral Evolution (SAVE) Program and through the National Institutes of Health Collaborative Influenza Vaccine Innovation Center (75N93021C00014) attributed to D.D.H. and the NIH, NIAID under contract number 75N93019C00051 attributed to A.G. We acknowledge funding support from the NSF (MCB-2032259) attributed to H.H.W. We thank all who contributed their data to GISAID. We express our gratitude to David Manthei, Carmen Gherasim, Victoria Blanc, Pamela Bennett-Baker, Savanna Sneeringer, Lauren Warsinske, Theresa Kowalski-Dobson, Alyssa Meyers, Zijin Chu, Hailey Kuiken, Lonnie Barnes, Ashley Eckard, Kathleen Lindsey, Dawson Davis, Aaron Rico, Gabriel Simjanovski, Mayurika Patel, and Nivea Vydiswaran of the IASO study team for supplying serum samples. We acknowledge Michael T. Yin and Magdalena E. Sobieszczyk at Columbia University Medical Center for providing serum samples.

## AUTHOR CONTRIBUTIONS

Lihong L. and D.D.H. conceived and supervised this project. Q.W. managed the project. Liyuan L., L.T.S., Y.H., Y.Q., and H.H.W. constructed the spike expression plasmids. Q.W., J.H., R.M.Z., and Lihong L. conducted pseudovirus neutralization assays. Q.W. and Lihong L. purified SARS-CoV-2 soluble spike proteins and hACE2 protein. Y.G. conducted bioinformatic analyses. Q.W., Lihong L., J.H., S.I., and J.Y. purified antibodies. Z.L. performed SPR assay. R.V., A.S.L., and A.G. provided clinical samples. Q.W., Y.G., Lihong L., and D.D.H. analyzed the results and wrote the manuscript. All authors reviewed the results and approved the final version of the manuscript.

## DECLARATION OF INTERESTS

Lihong L., S.I., J.Y., and D.D.H. are inventors on a provisional patent application on 10-40 described in this manuscript, titled “Isolation, characterization, and sequences of potent and broadly neutralizing monoclonal antibodies against SARS-CoV-2 and its variants as well as related coronaviruses” (63/271,627). D.D.H. is a co-founder of TaiMed Biologics and RenBio, consultant to WuXi Biologics and Brii Biosciences, and board director for Vicarious Surgical. Aubree Gordon serves on a scientific advisory board for Janssen Pharmaceuticals. Other authors declare no competing interests.

## MATERIALS & METHODS

### Human subjects

To evaluate neutralization sensitivity of BA.2.86 in this study, serum samples from three different clinical cohorts were utilized, which were “3 shots monovalent + 2 shots bivalent”, “BA.2 breakthrough” and “XBB breakthrough” cohorts. Sera of the first cohort were from healthy donors who had received three doses of SARS-CoV-2 monovalent mRNA vaccines (either Moderna mRNA-1273 or Pfizer BNT162b2), followed by two doses of bivalent mRNA vaccines. The latter two consisted of patients who had a BA.2 and a XBB breakthrough infection after multiple vaccinations, respectively.

Eight BA.2 breakthrough samples studied in this project were collected at Columbia University Irving Medical Center by Michael T. Yin’s and Magdalena E. Sobieszczyk’s teams. The remaining samples were collected at the University of Michigan through the Immunity-Associated with SARS-CoV-2 Study (IASO), which is an ongoing cohort study in Ann Arbor, Michigan that began in 2020^39^. All participants provided written informed consent and all serum samples were collected under protocols reviewed and approved by the Institutional Review Board of Columbia University or the Institutional Review Board of the University of Michigan Medical School.

IASO participants complete weekly symptom surveys and are tested for SARS-CoV-2 upon report of symptoms. All samples were examined by anti-nucleoprotein (NP) enzyme-linked immunoassay to confirm status of prior SARS-CoV-2 infection. Infected strains were confirmed by sequencing.

### Cell lines

HEK293T cells (CRL-3216) for pseudovirus producing and Vero-E6 cells (CRL-1586) for neutralization assays were purchased from the American Type Culture Collection. Expi293 cells (A14527) used for protein expression and purification, were purchased from Thermo Fisher Scientific. All cells were maintained according to the manufacturers’ instructions. The morphology of each cell line was confirmed visually before use. All cell lines tested negative for mycoplasma. Vero-E6 cells are from African green monkey kidneys. HEK293T cells and Expi293 cells are of female origin.

### Antibody and spike protein purification

To make antibodies and hACE2 with a Fc tag, the constructs encoding heavy and light chains of each antibody and the construct encoding hACE2 with Fc tag were/was transfected into Expi293 cells using 1 mg/mL polyethylenimine (PEI), respectively. Five days post transfection, cell supernatants were collected and clarified, and the expressed antibody and hACE2-Fc in cell supernatants were purified by using rProtein A Sepharose (GE).

The soluble spike constructs were transfected into Expi293 cells using 1 mg/mL polyethylenimine (PEI). Five days post transfection, cell supernatants were collected and clarified, and the spike proteins with a His tag were purified from the supernatant by using nickel-nitrilotriacetic acid (Ni-NTA) Sepharose (GE).

### Construction of SARS-CoV-2 spike plasmids

Spike-expressing plasmids for BA.2, XBB.1.5, and EG.5.1 pseudovirus generation were made in the previous studies^2,40^,41. Spike-expressing plasmids of BA.2.86-V1 and BA.2.86-V2 variants, as well as individual mutations found in BA.2.86 in the BA.2 background, were generated by MEGAA^42^ as previously described^4,10.^ Briefly, 5’-phosphorylated oligo pools with designed mutations were synthesized from SYNTAX Platform (Model STX-200) and Integrated DNA Technologies. The corresponding regions of the BA.2 spike gene construct were replaced with oligos by using annealing, extension, ligation, and PCR steps. To confirm the sequences of the variants, next generation sequencing^43^ and Oxford Nanopore sequencing were performed on the Illumina Miseq platform (single-end mode with 50 bp R1) and on a MinION with the MinKNOW v21.11.8 (Oxford Nanopore Technologies). Using Cutadapt v2.1^44^, Bowtie2 v2.3.4^45^, Guppy v3.6.0 (Oxford Nanopore Technologies) in GPU mode, and a custom Python script, we trimmed, aligned, basecalled, and filtered the reads for the full-length spike genes, respectively, and then viewed the read alignments in Integrative Genomics Viewer^46^.

To make soluble spike proteins of SARS-CoV-2 variants investigated in this study, we generated plasmid constructs encoding ectodomains (1-1208aa based on the sequence numbering of WA1) of spike proteins. In addition, these constructs also have a GSAS substitution at furin cleavage site (682-685aa which are RRAR in WA1) and a 2P substitution at positions K986 and V987 and are fused with a foldon tag followed by a 6× His tag. All constructs were confirmed by Sanger sequencing.

### Pseudovirus production

To make SARS-CoV-2 pseudoviruses, the spike-expressing plasmids were transfected into HEK293T cells using 1 mg/mL PEI. One day post transfection, cells were infected with VSV-G pseudotyped ΔG-luciferase (G*ΔG-luciferase, Kerafast) at a multiplicity (MOI) of ∼3 to 5. Two hours after infection, VSV-G pseudotyped ΔG-luciferase was removed by washing the cells with PBS three times. Cells were then maintained in fresh medium for another day before the cell supernatants containing pseudoviruses were harvested, clarified by centrifugation, aliquoted, and stored at -80°C.

### Pseudovirus neutralization assay

Before conducting neutralization assays, pseudoviruses were titrated on Vero-E6 cells to normalize the viral input between different viruses and assays. Serum samples were inactivated at 56 °C for 30 minutes before use. For serum neutralization assays, inactivated sera were diluted from 12.5-fold with a dilution factor of four. For mAb neutralization assays, mAbs were diluted from 20 μg/mL with a dilution factor of five. Dilutions were performed in 96 well plates in triplicates. Then 50 μL of each dilution of serum or mAb was incubated with 50 μL diluted pseudovirus for 1 hour at 37°C, followed by adding 100 μL of resuspended Vero-E6 cells at a density of 4 × 10^6^ cells/mL. Wells with no serum or no mAb (meaning virus alone) were included in all plates. Plates were then incubated at 37°C overnight before luciferase activity was quantified using the Luciferase Assay System (Promega) on SoftMax Pro v.7.0.2 (Molecular Devices). The reduction in luciferase activity for each serum and mAb dose, when compared with the “virus alone” controls, was calculated. Neutralization ID_50_ values for sera and IC_50_ values for mAbs were obtained by fitting a nonlinear five-parameter dose-response curve to the data in GraphPad Prism v.9.2.

### Phylogenetic analysis

Genome sequences of SARS-CoV-2 subvariants were retrieved from the GISAID (global initiative on sharing avian flu data) database. Subsequently, the spike protein sequences were extracted from the genomes using an in-house python script. The spike protein sequences were then aligned using the MUSCLE software (version 3.8.31). Low-quality sequencing sites characterized by the presence of ‘N’ were manually curated to ensure the mutations fit the consensus mutations in each variant. A Maximum Likelihood phylogenetic tree was constructed using MEGA11, utilizing the Tamura-Nei model and validated with 500 bootstrap replications.

### Antigenic cartography

Antigenic distances between sera to BA.2 and other SARS-CoV-2 variants were determined by integrating ID_50_ values of individual serum samples through a published antigenic cartography approach^47^. The visualization was generated using the Racmacs package (v.1.1.4, https://acorg.github.io/Racmacs/) in R version 4.0.3. With optimization steps set at 2,000 and the minimum column basis parameter set to ‘none’, the ‘mapDistances’ function was employed to calculate antigenic distances between each serum sample and variant. The final distances were represented by the average distances from all sera to each variant. BA.2 served as the center of sera for each group, the seeds for each antigenic map were manually adjusted to ensure that EG.5.1 was displayed in the horizontal direction relative to BA.2.

### Structural modeling

The structures of antibody–RBD complexes for modeling were obtained from PDB (PDB IDs: 7WKA for S3H3, 8D8Q for COV2-2130, 7MMO for LY-CoV1404, 7TAS for S2K146, and 7TCA for A19-46.1). The electrostatic potential was estimated by APBS electrostatics plugin, and the mutagenesis analysis were performed by Pymol version 2.5.4 (Schrödinger, LLC). The PDB ID for the BA.2 RBD and hACE2 complex is 7ZF7. The interaction residues of footprints were identified by PDBePISA^48^.

### Surface plasmon resonance (SPR)

The CM5 chip was immobilized with anti-His antibodies utilizing the His Capture Kit (Cytiva) to facilitate the capture of the spike protein via its C-terminal His-tag. Thereafter, a serial dilution of the hACE2 protein fused with a Fc tag was introduced over the chip, prepared in the HBS-EP+ buffer (Cytiva). Binding affinities were ascertained using the Biacore T200 system, operating at 25°C in a single-cycle mode. Subsequently, the acquired data were scrutinized using the Evaluation Software, adhering to a 1:1 binding model.

### QUANTIFICATION AND STATISTICAL ANALYSIS

IC_50_ and ID_50_ values were determined by fitting the data to five-parameter dose-response curves in GraphPad Prism v.9.2. Comparisons were made by two-tailed Wilcoxon matched-pairs signed-rank tests. ns, not significant; \**p* < 0.05, \**p* < 0.01; \*\*\**p* < 0.001; \*\*\*\**p* < 0.0001.

## Notes

### Summary of Updates

In the current version, Figures 3 and 4 are not displaying as intended. We've made corrections and will be uploading the updated figures.

