## supplemental data for "Antigenicity and receptor affinity of SARS-CoV-2 BA.2.86 spike"

### Extended Data Table 1. Demographics of clinical cohorts.

| Sample ID | Vaccine type and infected strain | Days post last vaccination or *infection | Documented COVID-19 | Age | Gender |
|---|---|---|---|---|---|
| 3 shots monovalent + 2 shots bivalent (n = 17) | | | | | |
| L1 | BNT162b2/BNT162b2/BNT162b2/BNT162b2/Pfizer Bivalent/Pfizer Bivalent | 33 | No | 66 | Female |
| L2 | BNT162b2/BNT162b2/BNT162b2/BNT162b2/Pfizer Bivalent/Pfizer Bivalent | 31 | No | 68 | Male |
| L3 | BNT162b2/BNT162b2/BNT162b2/BNT162b2/Pfizer Bivalent/Pfizer Bivalent | 28 | No | 74 | Male |
| L4 | BNT162b2/BNT162b2/BNT162b2/BNT162b2/Pfizer Bivalent/Pfizer Bivalent | 32 | No | 64 | Female |
| L5 | BNT162b2/BNT162b2/BNT162b2/BNT162b2/Pfizer Bivalent/Pfizer Bivalent | 33 | No | 64 | Female |
| L6 | BNT162b2/BNT162b2/BNT162b2/BNT162b2/Pfizer Bivalent/Pfizer Bivalent | 34 | No | 65 | Female |
| L7 | BNT162b2/BNT162b2/BNT162b2/BNT162b2/Pfizer Bivalent/Pfizer Bivalent | 29 | No | 70 | Male |
| L8 | mRNA-1273/mRNA-1273/mRNA-1273/mRNA-1273/Moderna Bivalent/Pfizer Bivalent | 33 | No | 63 | Male |
| L9 | BNT162b2/BNT162b2/BNT162b2/BNT162b2/Pfizer Bivalent/Pfizer Bivalent | 33 | No | 63 | Male |
| L10 | AstraZeneca/AstraZeneca/BNT162b2/BNT162b2/Pfizer Bivalent/Pfizer Bivalent | 43 | No | 73 | Male |
| L11 | BNT162b2/BNT162b2/BNT162b2/BNT162b2/Pfizer Bivalent/Pfizer Bivalent | 34 | No | 66 | Male |
| L12 | BNT162b2/BNT162b2/BNT162b2/BNT162b2/Pfizer Bivalent/Pfizer Bivalent | 26 | No | 78 | Male |
| L13 | BNT162b2/BNT162b2/BNT162b2/BNT162b2/Pfizer Bivalent/Pfizer Bivalent | 33 | No | 68 | Female |
| L15 | BNT162b2/BNT162b2/BNT162b2/BNT162b2/Pfizer Bivalent/Pfizer Bivalent | 31 | No | 74 | Male |
| L16 | BNT162b2/BNT162b2/BNT162b2/BNT162b2/Pfizer Bivalent/Pfizer Bivalent | 46 | No | 94 | Male |
| L17 | BNT162b2/BNT162b2/BNT162b2/BNT162b2/Moderna Bivalent/Pfizer Bivalent | 21 | No | 64 | Female |
| L18 | mRNA-1273/mRNA-1273/mRNA-1273/mRNA-1273/Moderna Bivalent/Moderna Bivalent | 42 | No | 60 | Female |
| BA.2 breakthrough (n = 25) | | | | | |
| Q35 | BNT162b2/BNT162b2/BA.2 | *14 | Yes | 50 | Female |
| Q36 | BNT162b2/BNT162b2/BNT162b2/Ad26.COV2.S/BA.2 | *22 | Yes | 69 | Male |
| Q50 | mRNA-1273/mRNA-1273/mRNA-1273/BA.2 | *14 | Yes | 34 | Male |
| Q51 | BNT162b2/BNT162b2/mRNA-1273/BA.2 | *19 | Yes | 33 | Female |
| Q52 | BNT162b2/BNT162b2/mRNA-1273/BA.2 | *18 | Yes | 29 | Female |
| C1 | BNT162b2/BNT162b2/BNT162b2/BA.2 | *28 | Yes | 22 | Male |
| C2 | mRNA-1273/mRNA-1273/mRNA-1273/BA.2 | *56 | Yes | 30 | Female |
| C3 | BNT162b2/BNT162b2/BNT162b2/BA.2 | *34 | Yes | 30 | Female |
| BA.2-10 | BNT162b2/BNT162b2/mRNA-1273/BA.2 | *30 | Yes | 59 | Female |
| BA.2-11 | BNT162b2/BNT162b2/BNT162b2/BA.2 | *29 | Yes | 39 | Female |
| BA.2-12 | BNT162b2/BNT162b2/BNT162b2/BA.2 | *18 | Yes | 45 | Female |
| BA.2-13 | BNT162b2/BNT162b2/BNT162b2/BNT162b2/BA.2 | *31 | Yes | 59 | Female |
| BA.2-14 | BNT162b2/BNT162b2/BNT162b2/BNT162b2/BA.2 | *25 | Yes | 39 | Male |
| BA.2-15 | BNT162b2/BNT162b2/BNT162b2/mRNA-1273/BA.2 | *32 | Yes | 61 | Male |
| BA.2-16 | Ad26.COV2.S/Ad26.COV2.S/BA.2 | *29 | Yes | 47 | Female |
| BA.2-17 | BNT162b2/BNT162b2/BNT162b2/BA.2 | *29 | Yes | 45 | Male |
| BA.2-18 | BNT162b2/BNT162b2/BNT162b2/mRNA-1273/BA.2 | *29 | Yes | 25 | Male |
| BA.2-19 | BNT162b2/BNT162b2/BNT162b2/BA.2 | *32 | Yes | 37 | Female |
| BA.2-20 | BNT162b2/BNT162b2/BNT162b2/BNT162b2/BA.2 | *31 | Yes | 62 | Male |
| BA.2-21 | BNT162b2/BNT162b2/BNT162b2/mRNA-1273/BNT162b2/BA.2 | *28 | Yes | 58 | Male |
| BA.2-22 | BNT162b2/BNT162b2/BNT162b2/BNT162b2/BA.2 | *30 | Yes | 29 | Male |
| BA.2-23 | BNT162b2/BNT162b2/BNT162b2/BA.2 | *28 | Yes | 57 | Male |
| BA.2-24 | mRNA-1273/mRNA-1273/BNT162b2/BA.2 | *29 | Yes | 47 | Male |
| BA.2-25 | BNT162b2/BNT162b2/BNT162b2/BA.2 | *31 | Yes | 47 | Male |
| BA.2-26 | Ad26.COV2.S/mRNA-1273/BA.2 | *31 | Yes | 46 | Male |
| XBB breakthrough (n =19) | | | | | |
| XBB-1 | BNT162b2/BNT162b2/BNT162b2/BNT162b2/BNT162b2/XBB | *35 | Yes | 38 | Female |
| XBB-2 | mRNA-1273/mRNA-1273/mRNA-1273/BNT162b2/Moderna bivalent/XBB | *50 | Yes | 38 | Male |
| XBB-3 | BNT162b2/BNT162b2/BNT162b2/Moderna bivalent/XBB | *54 | Yes | 61 | Male |
| XBB-4 | BNT162b2/BNT162b2/mRNA-1273/BNT162b2/XBB | *23 | Yes | 41 | Male |
| XBB-5 | BNT162b2/BNT162b2/BNT162b2/mRNA-1273/XBB | *14 | Yes | 37 | Male |
| XBB-6 | BNT162b2/BNT162b2/mRNA-1273/BNT162b2/XBB | *78 | Yes | 41 | Male |
| XBB-7 | BNT162b2/BNT162b2/BNT162b2/mRNA-1273/XBB | *79 | Yes | 37 | Male |
| XBB-8 | BNT162b2/BNT162b2/BNT162b2/BNT162b2/BNT162b2/XBB | *86 | Yes | 61 | Female |
| XBB-11 | BNT162b2/BNT162b2/BNT162b2/BNT162b2/XBB | *30 | Yes | 33 | Male |
| XBB-12 | BNT162b2/BNT162b2/BNT162b2/BNT162b2/BNT162b2/XBB | *21 | Yes | 52 | Male |
| XBB-13 | BNT162b2/BNT162b2/BNT162b2/XBB | *18 | Yes | 59 | Male |
| XBB-14 | BNT162b2/BNT162b2/BNT162b2/BNT162b2/BNT162b2/XBB | *32 | Yes | 50 | Female |
| XBB-15 | mRNA-1273/mRNA-1273/mRNA-1273/BNT162b2/XBB | *56 | Yes | 60 | Male |
| XBB-16 | mRNA-1273/mRNA-1273/mRNA-1273/mRNA-1273/BNT162b2/XBB | *24 | Yes | 59 | Male |
| XBB-17 | BNT162b2/BNT162b2/BNT162b2/BNT162b2/XBB | *28 | Yes | 40 | Male |
| XBB-18 | Ad26.COV2.S/Ad26.COV2.S/mRNA-1273/BNT162b2/XBB | *29 | Yes | 46 | Female |
| XBB-19 | BNT162b2/BNT162b2/BNT162b2/BNT162b2/BNT162b2/mRNA-1273/XBB | *19 | Yes | 76 | Female |
| XBB-20 | BNT162b2/BNT162b2/mRNA-1273/mRNA-1273/XBB | *28 | Yes | 61 | Female |
| XBB-21 | BNT162b2/BNT162b2/BNT162b2/BNT162b2/XBB | *35 | Yes | 45 | Male |

**Extended Data Table 2. Neutralization $IC_{50}$ values of BA.2, XBB.1.5, EG.5.1, BA.2.86-V1 and BA.2 carrying individual spike mutations found in BA.2.86 by mAbs.**

| $IC_{50}$ (μg/mL) | NTD | NTD-SD2 | SD1 | | | RBD class 1 | | | | | | | | RBD class 2 | | | | RBD class 3 | | | | | RBD class 4/1 | | |
|---|---|---|---|---|---|---|---|---|---|---|---|---|---|---|---|---|---|---|---|---|---|---|---|---|---|
| | C1520 | C1717 | S3H3 | ADARC1 | ADARC2 | S2K146 | BD57-0129 | BD56-1302 | BD56-1854 | Omi-3 | Omi-18 | BD-515 | Omi-42 | COV2-2196 | XGv347 | ZCB11 | XGv051 | A19-46.1 | S309 | COV2-2130 | LY-COV1404 | Beta-54 | BD55-4637 | SA55 | 10-40 |
| BA.2 | 0.001 | 0.451 | 0.016 | 0.087 | 0.063 | 0.039 | 0.003 | 0.003 | 0.001 | 0.011 | 0.003 | 0.012 | 0.023 | 0.123 | 0.001 | 0.004 | 0.001 | 0.045 | 0.494 | 0.006 | 0.001 | 0.146 | 0.023 | 0.010 | 7.515 |
| XBB.1.5 | >10 | 0.358 | 0.010 | 0.103 | 0.053 | 0.104 | 0.005 | 0.012 | 0.007 | 0.385 | 2.030 | 0.157 | 0.014 | >10 | >10 | >10 | 0.340 | >10 | 0.343 | >10 | >10 | 0.349 | 0.112 | 0.018 | 7.252 |
| EG.5.1 | >10 | 0.779 | 0.021 | 0.162 | 0.085 | 0.057 | 0.003 | 0.012 | 0.012 | >10 | >10 | >10 | 7.459 | >10 | >10 | >10 | 0.580 | >10 | 0.192 | >10 | >10 | 0.329 | 0.064 | 0.014 | 5.397 |
| BA.2.86-V1 | 0.215 | 0.693 | >10 | >10 | >10 | >10 | 0.003 | 0.006 | 0.002 | 0.450 | 0.522 | 0.115 | 0.017 | >10 | >10 | >10 | >10 | >10 | 9.130 | >10 | >10 | >10 | 0.019 | 0.001 | 1.714 |
| V16_insMPLF | 0.003 | 0.770 | 0.010 | 0.089 | 0.028 | 0.056 | 0.007 | 0.005 | 0.003 | 0.026 | 0.005 | 0.015 | 0.034 | 0.224 | 0.003 | 0.018 | 0.002 | 0.025 | 0.366 | 0.008 | 0.001 | 0.367 | 0.025 | 0.015 | 9.476 |
| BA.2-R21T | 0.002 | 0.350 | 0.014 | 0.084 | 0.074 | 0.033 | 0.004 | 0.003 | 0.002 | 0.020 | 0.004 | 0.012 | 0.031 | 0.180 | 0.002 | 0.016 | 0.001 | 0.043 | 0.430 | 0.006 | 0.001 | 0.267 | 0.037 | 0.014 | 4.374 |
| BA.2-S50L | 0.001 | 0.185 | 0.018 | 0.038 | 0.099 | 0.007 | 0.002 | 0.002 | 0.002 | 0.006 | 0.002 | 0.004 | 0.011 | 0.103 | <0.001 | <0.001 | <0.001 | 0.008 | 0.171 | 0.003 | 0.001 | 0.052 | 0.010 | 0.003 | 1.060 |
| BA.2-H60/V70Δ | 0.001 | 0.505 | 0.015 | 0.073 | 0.061 | 0.036 | 0.004 | 0.003 | 0.003 | 0.013 | 0.003 | 0.006 | 0.027 | 0.170 | 0.001 | 0.008 | <0.001 | 0.033 | 0.408 | 0.006 | 0.001 | 0.159 | 0.019 | 0.009 | 9.432 |
| BA.2-V127F | 0.001 | 0.499 | 0.010 | 0.062 | 0.156 | 0.026 | 0.005 | 0.003 | 0.002 | 0.015 | 0.004 | 0.009 | 0.027 | 0.252 | 0.002 | 0.008 | 0.002 | 0.047 | 0.457 | 0.008 | <0.001 | 0.248 | 0.028 | 0.012 | 8.461 |
| BA.2-Y144Δ | 0.001 | 0.362 | 0.012 | 0.066 | 0.051 | 0.026 | 0.007 | 0.006 | 0.001 | 0.020 | 0.006 | 0.013 | 0.029 | 0.236 | 0.003 | 0.012 | 0.002 | 0.024 | 0.408 | 0.008 | 0.001 | 0.221 | 0.031 | 0.015 | 9.400 |
| BA.2-F157S | 0.001 | 0.381 | 0.010 | 0.097 | 0.051 | 0.029 | 0.003 | 0.003 | 0.002 | 0.014 | 0.004 | 0.008 | 0.025 | 0.119 | 0.001 | 0.003 | 0.001 | 0.030 | 0.307 | 0.004 | 0.001 | 0.203 | 0.018 | 0.009 | 6.954 |
| BA.2-R158G | 0.001 | 0.257 | 0.008 | 0.080 | 0.047 | 0.033 | 0.008 | 0.004 | 0.002 | 0.012 | 0.003 | 0.006 | 0.016 | 0.163 | 0.001 | 0.008 | <0.001 | 0.030 | 0.281 | 0.007 | 0.001 | 0.280 | 0.017 | 0.013 | 6.318 |
| BA.2-N211Δ | 0.001 | 0.171 | 0.009 | 0.093 | 0.054 | 0.015 | 0.003 | 0.003 | 0.002 | 0.014 | 0.003 | 0.008 | 0.030 | 0.105 | 0.001 | 0.004 | <0.001 | 0.030 | 0.235 | 0.005 | 0.001 | 0.206 | 0.013 | 0.009 | 3.509 |
| BA.2-L212I | 0.001 | 0.486 | 0.012 | 0.069 | 0.055 | 0.044 | 0.004 | 0.003 | 0.002 | 0.015 | 0.002 | 0.007 | 0.022 | 0.195 | 0.001 | 0.007 | <0.001 | 0.043 | 0.210 | 0.006 | 0.001 | 0.238 | 0.016 | 0.013 | 5.188 |
| BA.2-L216F | 0.001 | 0.696 | 0.016 | 0.109 | 0.054 | 0.023 | 0.003 | 0.002 | 0.001 | 0.012 | 0.003 | 0.007 | 0.020 | 0.186 | 0.002 | 0.009 | 0.001 | 0.039 | 0.388 | 0.005 | 0.001 | 0.181 | 0.015 | 0.007 | 5.708 |
| BA.2-H245N | 0.160 | 0.484 | 0.013 | 0.105 | 0.160 | 0.042 | 0.005 | 0.004 | 0.002 | 0.015 | 0.004 | 0.010 | 0.024 | 0.185 | 0.002 | 0.009 | 0.002 | 0.044 | 0.503 | 0.008 | <0.001 | 0.241 | 0.024 | 0.011 | 6.877 |
| BA.2-A264D | 0.001 | 0.338 | 0.012 | 0.081 | 0.052 | 0.041 | 0.003 | 0.003 | 0.001 | 0.017 | 0.004 | 0.009 | 0.018 | 0.161 | 0.002 | 0.007 | 0.001 | 0.030 | 0.378 | 0.008 | <0.001 | 0.236 | 0.020 | 0.009 | 5.380 |
| BA.2-I332V | 0.001 | 0.361 | 0.017 | 0.100 | 0.217 | 0.014 | 0.002 | 0.001 | 0.001 | 0.005 | 0.002 | 0.005 | 0.010 | 0.100 | <0.001 | <0.001 | 0.001 | 0.016 | 0.913 | 0.004 | 0.001 | 0.102 | 0.013 | 0.004 | 1.431 |
| BA.2-D339H | 0.001 | 0.739 | 0.015 | 0.061 | 0.038 | 0.038 | 0.006 | 0.004 | 0.002 | 0.018 | 0.005 | 0.011 | 0.024 | 0.198 | 0.001 | 0.010 | 0.001 | 0.036 | 0.108 | 0.004 | 0.001 | 0.426 | 0.024 | 0.014 | 7.557 |
| BA.2-K356T | 0.001 | 0.471 | 0.006 | 0.070 | 0.030 | 0.049 | 0.009 | 0.005 | 0.003 | 0.037 | 0.005 | 0.013 | 0.036 | 0.370 | 0.001 | 0.014 | 0.001 | 0.338 | >10 | 0.007 | 0.001 | 0.114 | 0.063 | 0.025 | 9.658 |
| BA.2-R403K | 0.002 | 0.172 | 0.019 | 0.087 | 0.315 | 0.011 | 0.002 | 0.002 | 0.001 | 0.004 | 0.002 | 0.003 | 0.008 | 0.097 | <0.001 | 0.002 | <0.001 | 0.027 | 0.244 | 0.005 | 0.001 | 0.071 | 0.006 | 0.002 | 1.678 |
| BA.2-V445H | 0.001 | 0.266 | 0.011 | 0.058 | 0.043 | 0.029 | 0.003 | 0.003 | 0.001 | 0.013 | 0.005 | 0.009 | 0.022 | 0.137 | 0.002 | 0.014 | 0.001 | 0.042 | 0.416 | 0.005 | >10 | >10 | 0.037 | 0.003 | 7.614 |
| BA.2-G446S | 0.001 | 0.328 | 0.007 | 0.071 | 0.041 | 0.016 | 0.003 | 0.002 | 0.001 | 0.010 | 0.003 | 0.008 | 0.018 | 0.238 | 0.001 | 0.006 | 0.001 | 0.065 | 0.332 | 0.007 | 0.001 | 0.011 | 0.028 | 0.006 | 4.425 |
| BA.2-N450D | 0.001 | 0.742 | 0.012 | 0.084 | 0.056 | 0.020 | 0.004 | 0.003 | 0.002 | 0.013 | 0.004 | 0.009 | 0.022 | 0.125 | 0.001 | 0.007 | 0.001 | 0.239 | 0.316 | 1.535 | 0.001 | 0.113 | 0.020 | 0.011 | 8.862 |
| BA.2-L452W | 0.001 | 0.611 | 0.015 | 0.064 | 0.068 | 0.032 | 0.004 | 0.003 | 0.002 | 0.019 | 0.003 | 0.007 | 0.019 | 0.191 | 0.001 | 0.008 | 0.002 | >10 | 0.614 | 0.009 | 0.001 | 0.383 | 0.030 | 0.012 | 7.834 |
| BA.2-N460K | 0.001 | 0.414 | 0.006 | 0.113 | 0.035 | 0.074 | 0.009 | 0.010 | 0.004 | 0.378 | 0.015 | 0.104 | 0.071 | 0.082 | <0.001 | 0.015 | 0.002 | 0.063 | 0.292 | 0.007 | 0.001 | 0.368 | 0.046 | 0.015 | 9.922 |
| BA.2-N481K | 0.002 | 0.448 | 0.014 | 0.066 | 0.086 | 0.026 | 0.005 | 0.003 | 0.002 | 0.015 | 0.003 | 0.008 | 0.022 | 0.327 | 0.003 | 0.011 | 0.001 | 0.050 | 0.301 | 0.008 | 0.001 | 0.206 | 0.017 | 0.007 | 7.480 |
| BA.2-V483Δ | 0.002 | 0.506 | 0.012 | 0.072 | 0.072 | 0.032 | 0.004 | 0.002 | 0.002 | 0.018 | 0.003 | 0.009 | 0.023 | 0.183 | 0.003 | 0.006 | 0.001 | 0.072 | 0.442 | 0.006 | 0.001 | 0.162 | 0.028 | 0.012 | 7.788 |
| BA.2-A484K | 0.001 | 0.424 | 0.009 | 0.064 | 0.053 | 0.399 | 0.005 | 0.004 | 0.002 | 0.015 | 0.002 | 0.012 | 0.033 | 0.293 | 0.002 | 0.006 | 0.001 | 0.740 | 0.404 | 0.007 | 0.001 | 0.119 | 0.021 | 0.009 | 7.142 |
| BA.2-F486P | 0.001 | 0.373 | 0.010 | 0.051 | 0.050 | >10 | 0.005 | 0.007 | 0.005 | 0.293 | 0.008 | 0.017 | 0.042 | >10 | >10 | >10 | 0.294 | 0.036 | 0.290 | 0.006 | 0.001 | 0.207 | 0.034 | 0.010 | 9.432 |
| BA.2-R493Q | 0.003 | 0.196 | 0.015 | 0.078 | 0.059 | 0.003 | 0.001 | 0.001 | 0.001 | 0.007 | 0.004 | 0.003 | 0.004 | 0.005 | <0.001 | <0.001 | <0.001 | 0.020 | 0.697 | 0.005 | 0.001 | 0.104 | 0.019 | 0.005 | 4.479 |
| BA.2-E554K | 0.001 | 0.428 | >10 | >10 | 0.374 | 0.027 | 0.004 | 0.004 | 0.002 | 0.013 | 0.003 | 0.010 | 0.015 | 0.133 | 0.001 | 0.007 | 0.001 | 0.031 | 0.289 | 0.008 | <0.001 | 0.225 | 0.016 | 0.010 | 5.629 |
| BA.2-A570V | 0.001 | 0.192 | 0.010 | 0.052 | 0.034 | 0.027 | 0.004 | 0.003 | 0.001 | 0.012 | 0.004 | 0.008 | 0.016 | 0.171 | 0.002 | 0.006 | 0.001 | 0.031 | 0.337 | 0.005 | <0.001 | 0.247 | 0.021 | 0.009 | 6.108 |
| BA.2-P621S | 0.001 | 0.410 | 0.058 | 0.112 | 0.040 | 0.022 | 0.003 | 0.002 | 0.001 | 0.013 | 0.002 | 0.007 | 0.012 | 0.313 | 0.002 | 0.006 | 0.001 | 0.035 | 0.234 | 0.004 | <0.001 | 0.118 | 0.016 | 0.013 | 2.849 |
| BA.2-I670V | 0.001 | 0.303 | 0.009 | 0.049 | 0.035 | 0.035 | 0.005 | 0.003 | 0.002 | 0.015 | 0.004 | 0.008 | 0.021 | 0.155 | 0.002 | 0.007 | 0.001 | 0.037 | 0.327 | 0.007 | 0.001 | 0.234 | 0.024 | 0.009 | 6.229 |
| BA.2-H681R | 0.001 | 0.311 | 0.008 | 0.063 | 0.040 | 0.018 | 0.004 | 0.003 | 0.001 | 0.015 | 0.004 | 0.007 | 0.018 | 0.137 | 0.001 | 0.007 | 0.001 | 0.041 | 0.394 | 0.007 | 0.001 | 0.221 | 0.023 | 0.012 | 8.412 |
| BA.2-S939F | 0.001 | 0.482 | 0.008 | 0.068 | 0.053 | 0.016 | 0.002 | 0.002 | 0.001 | 0.011 | 0.003 | 0.005 | 0.015 | 0.162 | 0.001 | 0.003 | 0.001 | 0.019 | 0.294 | 0.006 | 0.001 | 0.175 | 0.016 | 0.008 | 2.737 |
| BA.2-P1143L | 0.001 | 0.341 | 0.010 | 0.049 | 0.037 | 0.014 | 0.004 | 0.003 | 0.001 | 0.011 | 0.003 | 0.005 | 0.015 | 0.101 | 0.001 | 0.003 | 0.001 | 0.021 | 0.228 | 0.007 | 0.001 | 0.267 | 0.019 | 0.010 | 4.796 |

| | | | | |
|---|---|---|---|---|
| >10 | <10 | <1 | <0.1 | <0.01 |

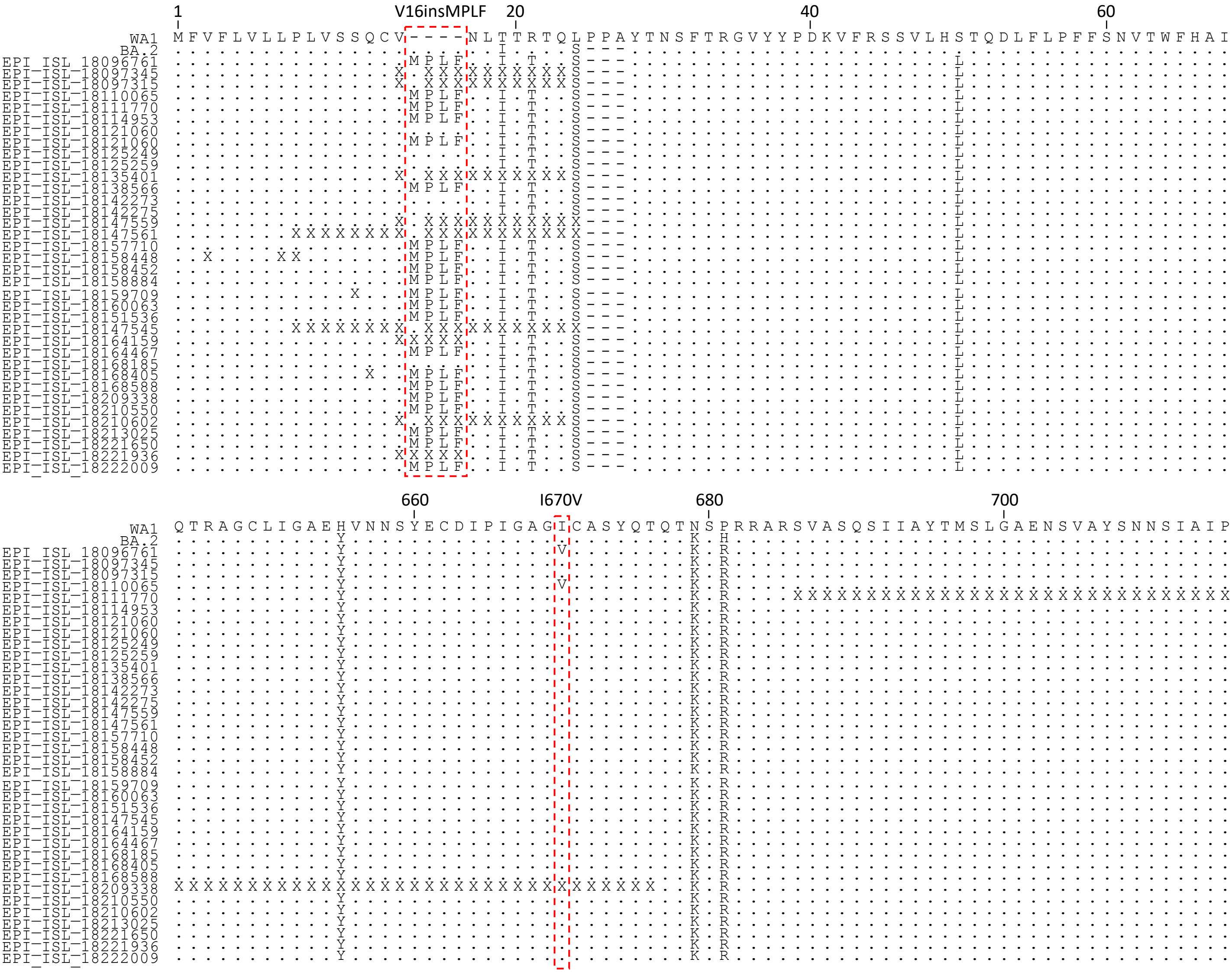


**Extended Data Figure 1. Spike sequence alignment of WA1 and BA.2 with BA.2.86 from human cases deposited to GISAID as of September 5, 2023.** The sequence numbering is based on WA1. Red boxes indicate the alignments of amino acids at position 16 and 670. “X”, low-quality sequencing data.
